# Model-based analysis of experimental hut data elucidates multifaceted effects of a volatile chemical on *Aedes aegypti* mosquitoes

**DOI:** 10.1101/164293

**Authors:** Quirine A. ten Bosch, Fanny Castro-Llanos, Hortance Manda, Amy C. Morrison, John P. Grieco, Nicole L. Achee, T.Alex Perkins

**Author notes:** Current address: Mathematical Modelling of Infectious Disease Unit, Institut Pasteur,Paris, France (Quirine A. ten Bosch).

## Abstract

**Background:** Insecticides used against *Aedes aegypti* and other disease vectors can elicit a multitude of dose-dependent effects on behavioral and bionomic traits. Estimating the potential epidemiological impact of a product requires thorough understanding of these effects and their interplay at different dosages. Volatile spatial repellent (SR) products come with an additional layer of complexity due to the potential for movement of affected mosquitoes or volatile particles of the product beyond the treated house. Here, we propose a statistical inference framework for estimating these nuanced effects of volatile SRs.

**Methods:** We fitted a continuous-time Markov chain model in a Bayesian framework to mark-release-recapture (MRR) data from an experimental hut study conducted in Iquitos, Peru. We estimated the effects of two dosages of transfluthrin on *Ae. aegypti* behaviors associated with human-vector contact: repellency, exiting, and knockdown in the treated space and in “downstream” adjacent huts. We validated the framework using simulated data.

**Results:** The odds of a female *Ae. aegypti* being repelled from a treated hut (*H*_T_) increased at both dosages (low dosage: odds = 1.64, 95% highest density interval (HDI) = 1.30-2.09; high dosage: odds = 1.35, HDI = 1.04-1.67). The relative risk of exiting from the treated hut was reduced (low: RR = 0.70, HDI = 0.62-1.09; high: RR = 0.70, HDI = 0.40-1.06), with this effect carrying over to untreated spaces as far as two huts away from the treated hut (*H*_2_) (low: RR = 0.79, HDI = 0.59-1.01; high: RR = 0.66, HDI = 0.50-0.87). Knockdown rates were increased in both treated and downstream huts, particularly under high dosage (*H*_T_: RR = 8.37, HDI = 2.11-17.35; *H*_1_: RR = 1.39, HDI = 0.52-2.69; *H*_2_: RR = 2.22, HDI = 0.96-3.86).

**Conclusions:** Our statistical inference framework is effective at elucidating multiple effects of volatile chemicals used in SR products, as well as their downstream effects. This framework provides a powerful tool for early selection of candidate SR product formulations worth advancing to costlier epidemiological trials, which are ultimately necessary for proof of concept of public health value and subsequent formal endorsement by health authorities.

## BACKGROUND

Insecticidal strategies against adult mosquitoes have been used extensively in the control of mosquito-borne diseases [1]. However, certain mosquito behaviors, such as outdoor and daytime biting, challenge the efficacy of traditional control tools like insecticide treated nets (ITNs) and indoor residual spraying (IRS) [2]. The evolution of physiological resistance to insecticides [3] and behavioral adaptation of mosquitoes [4, 5] also pose limitations to the effectiveness of such products.

The effect of vector control products often goes beyond their acute lethal effects. For example, ITNs can elicit knockdown with potential for mosquito recovery and can divert mosquitoes away from a protected human to alternate hosts [6-8]. Volatile chemicals such as transfluthrin and metofluthrin can be delivered in high dosages and result in high lethality, but they can likewise be formulated at lower dosages where acute toxicity is attenuated and other mosquito behaviors are elicited instead, as was described previously for residual pyrethroids [9, 10]. Currently, the term “spatial repellency” is used to describe a range of behaviors that products with volatile chemicals – including spatial repellents (SR) – may invoke [11], including repellency or reduced entry, irritancy or increase in exiting, and reduced biting [12-14]. These modes of action can have a concerted impact on disease transmission on an individual and community level [15-18].

Mark-release-recapture (MRR) experiments in natural settings, in which mosquitoes are marked with a unique color according to the location where they are released, offer unique opportunities to elucidate downstream dosage and behavioral effects of SRs by measuring lethality, repellency, and irritancy of a target vector species [10, 19-22].However, studies such as these have not yet provided the granularity required to disentangle distinct behavioral and bionomic effects. The primary challenge associated with the design and interpretation of these studies is that each mosquito is only observed once: when knocked down or when trapped in entry or exit traps. This leaves movement trajectories in between release and recapture locations unobserved, making it challenging to quantify the relative contributions of multiple competing effects that could account for observed individual-level outcomes under a multitude of equally plausible scenarios. One recent study [23] showed that even short periods of transient exposure to volatile SRs can have significant, and sometimes delayed, effects on vectors. Such unobserved effects may compromise traditional statistical analyses.

Models used for the analysis of MRR data have a long history in ecology [24-28].Originally developed to estimate survival probabilities and population sizes [29], they are now increasingly being used to inform spatial processes [30]. These models partition animal movement trajectories into states (e.g., breeding or foraging), with multi-state MRR models accounting for the probability of the animal occupying any of the possible states at a given time. Given sufficient information from sampling at multiple points in time and appropriate model constraints, these models can be extended for parameter estimation in the presence of unobserved states [31]. Bayesian methods are increasingly being applied to these types of problems given their treatment of all quantities as random variables [32-34]. These methods allow for formal treatment and quantification of parameter uncertainty, and they allow researchers to explicitly build on previous studies.

Here, we make a major advance in the technical capability to infer effects of SR products on adult female *Ae. aegypti* by developing a hierarchical Bayesian model and applying it to an MRR study conducted in Iquitos, Peru. We first demonstrate the accuracy of this approach using data simulated under the same design as in our field experiments. We then demonstrate the dose-dependency of knockdown, repellency, and exiting effects of the SR in both treated and untreated huts. We discuss the potential use of this framework to inform the projected impact and implementation of SRs and other vector control tools with volatile chemicals.

## METHODS

### Product

Technical grade transfluthriun (Sigma), a volatile pyrethroid insectide, was applied to cotton strips at 1/16^th^ and 1/8^th^ dilutions of the field application rate (FAR) (0.04g/m^2^) using previously established protocols [35]. Control strips of matched cotton material were treated with acetone alone. Cotton material was applied to the interior walls of the huts using magnets and metal frames [35].

### Experimental huts

A unique experimental hut configuration was used in which five independent structures were positioned adjacent to one another in a single row creating adjoining walls. Eave gaps were open, allowing a continuum of indoor space available for mosquito and volatile chemical movement throughout all five huts regardless of where the mosquitoes were released or where the product was applied (Figure 1). This design mimicked housing configurations common to the study location in Iquitos, Peru and reflects housing common to other dengue-endemic areas (i.e., urban settings in resource-poor, tropical areas). The huts measured 4 m wide x 6 m long and had 2 m high sidewalls. Each hut had two windows (one each on the front and back walls) equipped with exit traps, and each hut had two doors (one of each on the front and back walls) equipped with an “upper” and “lower” trap. The two outermost huts had additional eave traps. Hut construction materials and structural design were based on previous MRR hut studies [19, 20]. The study was performed at the Instituto Veterinario de Investigaciones in Iquitos, Peru (73.2 °W, 7.3 °S).

**Figure 1:**
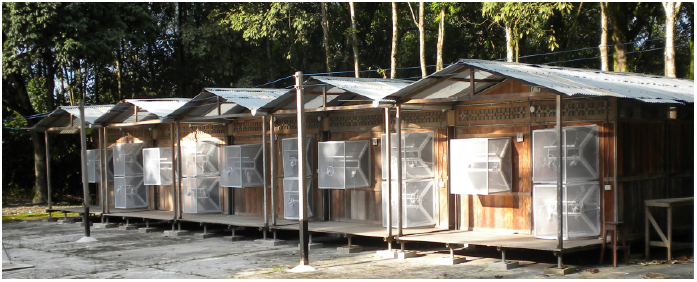
Experimental huts in row house configuration. Iquitos, Peru.

### Mosquitoes

Female *Ae. aegypti* test populations (F_1_-F_2_ generations) of 5-7 days old were reared from field-collected larvae following previously established protocols [36]. Mosquitoes were not blood-fed but were provided with cotton soaked with sucrose solution until 24 hours before being released in the experimental huts. Prior to release, five cohorts of 25 female mosquitoes each were marked with a unique color of fluorescent powder that corresponded to a single, specific experimental hut in which a cohort was released.

### Study design

The study was performed using previously described collection protocols [35, 37]. Transfluthrin-treated cotton was applied to the walls of the center hut (*H*_T_), while solvent-only material (control) was applied to the remaining huts. This design was intended to reflect a scenario whereby an SR product is only used by one homeowner in a group of houses. In each of the huts (*H*_2L_, *H*_1L_, *H*_T_, *H*_1R_, *H*_2R_) there was a human present under an untreated bed net to generate host-seeking cues and to monitor indoor knockdown. On each experimental day, mosquito test cohorts were released inside each of the untreated huts at 0530 hours. Mosquitoes within exit-traps were captured by two-person outdoor collection teams (five teams total) every 30 minutes from 0600 until 1800 hours.Collector teams rotated among huts at each sampling period to control for observer bias. Knockdown was monitored every hour by indoor collectors. Indoor collectors were rotated among huts at the end of a single day’s experiment to limit host cue bias. At 1800 hours, hand-held Prokopack aspirators [38] were used to recapture any remaining mosquitoes inside each hut that were not knocked down or had not exited and to calculate loss to follow-up. Color codes were used to record release origin and location of recapture in a single day. All recaptured mosquitoes (those from aspiration and in traps) were held with access to sugar source to monitor 24 hr mortality. Three trials were performed: 1) baseline (no chemical application), 2) transfluthrin at 1/8 FAR, and 3) transfluthrin at 1/16 FAR. A single trial consisted of five experimental days (i.e., five replicates). Movement during the baseline trial was measured prior to transfluthrin-integrated trials to monitor residual impact of treatment across trials.

### Model

*Continuous-time Markov chain* – A continuous-time Markov chain model was developed for the analysis of these data [39]. At any given time, mosquitoes can occupy any one of five huts (transient states: H_2L_, H_1L_, H_T_, H_1R_, or H_2R_) or have experienced one of 15 events represented by the absorbing states: X_2L_, X_1L_, X_T_, X_1R_, or X_2R_ for the exit traps in each hut, K_2L_, K_1L_, K_T_, K_1R_, or K_2R_ for knockdown in each hut, and U_2L_, U_1L_, U_T_, U_1R_, or U_2R_ for mosquitoes that were unaccounted for at the end of the experiment and were thus lost to follow-up at some unknown time. The infinitesimal generator matrix **A** contains the rates at which mosquitoes leave one state to move to the next, such that a_ij_ gives the rate at which a mosquito in state i moves to state j. These rates were assumed to be independent of time or previous trajectories; therefore, the time spent in state i before leaving follows an exponential distribution with mean 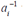 with 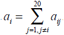 Note that the rates out of the absorbing states are zero and that, given symmetry in the system, the rates for hut 2L and 2R are equivalent (likewise for 1L and 1R). Subscripts in **A** indicate the distance from the treatment hut. The 20x20 matrix **A** is defined as

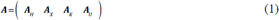

with

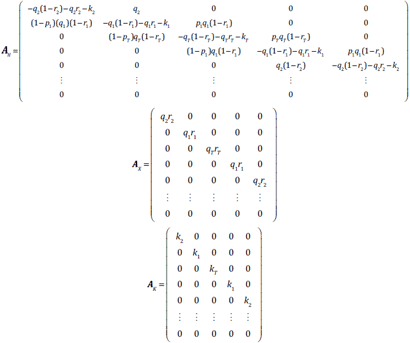

and

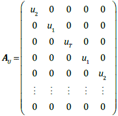

The rate *q*_*i*_ signifies the movement rate out of a hut. The direction of this movement depends on *r*_i_ (proportion of movement directed to outdoors) and, for *H*_1_, it further depends on repellency *p*_1_ (defined as the proportion of indoor movement directed away from *H*_T_). The knockdown rate *k*_*i*_ is allowed to vary by hut, whereas the loss to follow-up rate *u* is assumed to be the same across huts. Hereafter, we refer to the exit rate *q*_i_*r*_i_ as *x*_i_ (Figure 2).

**Figure 2:**
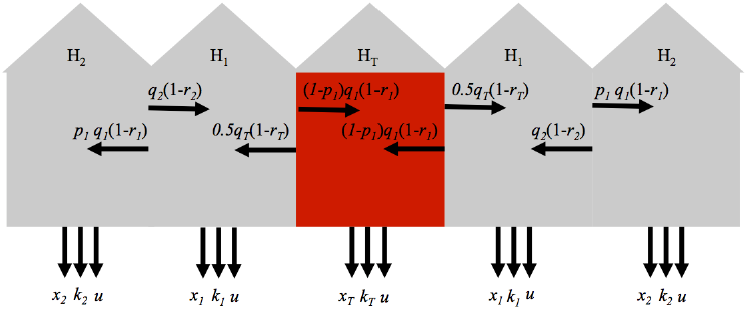
Illustration of experimental hut design and associated model parameters, with *q* = movement rate, *p* = proportion of between-hut movement directed away from the treated hut (repellency), *r* = proportion of movement directed outdoors, *q r* = *x* = exit rate, *k* = knockdown rate, and *u* = loss to follow-up rate. The red hut is the treated hut H_T_ where the SR treatment is applied. The subscripts indicate whether the parameter applies to H_T_ (subscript *T*) or to a hut one or two removed from H_T_.

The dynamics of the probabilities ***P***_ij_(*t*) of occupying any of the 20 states are governed by a system of differential equations with rates ***A*** and are known as the backward Kolmogorov differential equations [39]

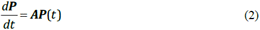

From this, we can derive the rates of change in the probability of occupying a given state

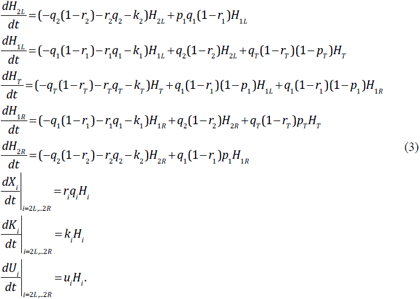

By initializing this system in one hut (e.g., *H*_2L_=1 and all other states are zero at *t*=0), solving for this system of differential equations gives the probability that a mosquito released in a given hut occupies a specific state at time *t*.

The absorbing states (i.e., *X*_i_*, K*_i_, and *U*_i_) represent competing endpoints in the sense that an individual who enters one of these states is no longer capable of entering any of the other states at some future time. The Markov chain accounts for competing endpoints vis-à-vis the property that the states are discrete and mutually exclusive. In addition, a mosquito released in 2L can only be knocked down in 2R conditional on having moved there prior to the knockdown event. The absence of non-zero rates to any of the absorbing states from other huts ensures this conditionality.

### Likelihoods

To estimate ***A***, we fitted eqn. (3) to the data using a likelihood-based approach. The data collected during the experiments consisted of a set of interval-and right-censored time-to-event data. Outcome measures of interest included exiting (i.e., leaving a space), knockdown, diversion (defined as the movement to a hut other than the release hut), and loss to follow-up (ltfu), where exiting, knockdown, and loss to follow-up are competing events. The cumulative, conditional probabilities for all events observed in the experiment can be directly obtained from the solutions of eqn. (3), as detailed in eqns. (4), (5), and (6).

*Interval-censored events*. – Data pertaining to knockdown and exit events are interval-censored between time points *t*_1_ and *t*_2_, with exit events recorded at 30-minute intervals and knockdown events at hourly intervals. Given model parameter set θ, the probability that a mosquito released in *H*_*rel*_ is observed to be knocked down in hut *H* at time *t*_2_ is

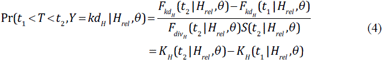

where *F*(*t*) denotes the probability that a specific event (here knockdown and movement to *H*) occurred in hut *H* by time *t* and *S*(*t*) denotes the survival function (i.e., the probability that no knockdown, exit, or loss to follow-up has occurred by time *t*). Exit and knockdown events contain indirect information on the diversion event, namely that the mosquito has moved from its release location to the hut where the event took place before the event occurred. This condition, as illustrated by F_div_ in the denominator of eqn. (4), is implicitly accounted for within eqn. (3); hence, the absence of conditioning in the second part of eqn. (4).

*Loss to follow-up. –* Of mosquitoes that are not retrieved at the end of the experiment, we know that they must have been lost to follow-up at some point between the start and the end of the experiment with probability

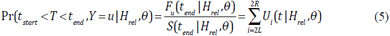

*Right-censored data*. – Mosquitoes retrieved by the end of the experiment are treated as right censored. Namely, the time before knockdown, exit, or ltfu would have occurred is longer than the duration of the study, but by how much is uncertain. In addition, we know that the mosquito moved from the release hut to the hut where it was retrieved with probability

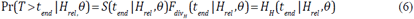

*Likelihood function. –* The overall likelihood of the parameters given the data is equal to the product of the probabilities of each individual observation conditional on the parameters. These observations include the number of mosquitoes exited or knocked down during specific time intervals during an experiment for different release huts, event huts, and experimental day, as well as numbers recaptured or lost to follow-up at the end of the experiment, resulting in

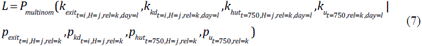

with *i* time points, *j* event-huts (*H*_2L_,*H*_2L_,*H*_T_,*H*_1L_,*H*_2R_,), *k* release-huts, and *l* experiment days (1 to 5). Each *k*_exit_, for instance, denotes the number of mosquitoes exiting from hut *H* observed at time *t*, by release hut and experiment day. The corresponding probabilities *p* are derived as detailed in eqns. (4), (5), and (6) and are assumed to be independent of the experiment day.

### Model fitting

We used a Bayesian Markov chain Monte Carlo (MCMC) approach for parameter estimation. Using Bayes’ theorem, we define the posterior probability density of the model’s parameters (θ) given the data as

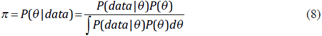

where P(θ) is the prior probability of the parameters. We utilized beta-distributed priors with median 0.5 for *p*_1_ and median 0.25 for *r*_*i*_ (i.e., a mosquito is three times as likely to move to an adjacent hut than to move outside), a gamma-distributed prior with mean 0.02 on the movement rates *q*_i_ (i.e., average time before moving to another hut of 50 minutes), and uniform priors for the remaining parameters (see Table 1 for distribution parameters). Average times before exiting from each hut (1/*q*_*i*_) were constrained between 5 minutes and 20 hours, and the average time until knockdown (1/*k*_*i*_) between 12 hours and 10 days [10, 19]. We explored the parameter space of θ more broadly using the Metropolis-Hastings algorithm.

**Table 1:**
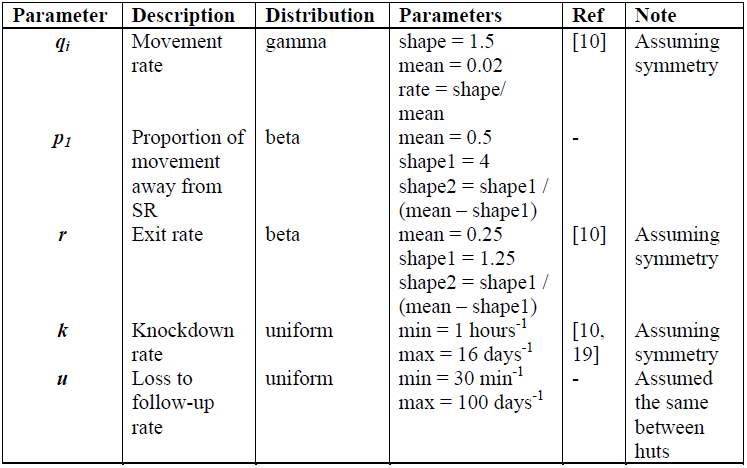
Parameter definitions and prior probability distributions for each.

We started from an initial parameter set θ_1_, which was randomly sampled from uniform distributions with bounds: *q*: 360^-1^-30^-1^, *p*_1_: 0.5-1, *r*_i_: 0-0.5, *k*: 1400^-1^-720^-1^, and *u*: 2000^-1^-1000^-1^. A new parameter was proposed such that λ_2_ = λ_1_+∧, where ∧ is a random value from a truncated normal proposal distribution *g* with mean λ_1_ and standard deviation formulated relative to λ_1_ and selected so as to ideally have an acceptance rate between 10% and 50% [40]. Which parameter was updated at a given iteration was determined by taking a random draw from a multinomial distribution with 11 categories (i.e., the number of model parameters to be estimated) and equal probabilities for each parameter. The probability for λ_2_ to be accepted depends on the likelihood of both θ_1_ and θ_2_ according to the Metropolis-Hastings rule as

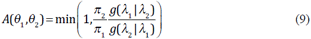

where θ_2_ differs from θ_1_ only with respect to λ and *g* denotes truncated normal proposal distributions (between zero and one for each *p* and *r*, and from zero to infinity otherwise):

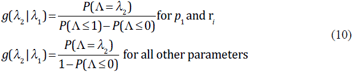

where ∧ is normally distributed with mean λ_1_ and standard deviations corresponding to each parameter’s proposal distribution.

In the event that the acceptance probability was larger than a randomly generated uniform value between zero and one, θ_2_ was accepted into the chain. Otherwise, θ_1_ was retained. Multiple iterations of this routine were performed (n = 90,000). This process was repeated five times starting from different initial parameter sets to assess convergence using the Gelman-Rubin (GR) statistic [41]. The resulting chains of accepted parameters (φ), after discarding a burn-in period (10,000), were combined to represent our sample from the posterior distribution (π).

### Simulation experiments

To validate the accuracy of the model-fitting algorithm, we simulated data with a known data-generating process corresponding to our likelihood formulation and with known model parameters. Probabilities for released mosquitoes to occupy a specific state over time were derived using eqn. (3). As follows from eqns. (4), these probabilities are defined for interval-and right-censored events. Random draws from a multinomial distribution with the simulated probabilities and a given number of released mosquitoes were taken to simulate numbers of mosquitoes occupying each state at the time points at which sampling was simulated to occur. In general, these simulation experiments were designed to mimic features of the empirical experiments.

Ten distinct simulated parameter sets were used to validate the accuracy of our statistical inference framework. These parameter sets were sampled from across the composite parameter space θ using the Sobol algorithm [42, 43], where the same bounds to this sampling space were applied as for the prior distributions (Table 1). Data were simulated for different numbers of released mosquitoes (25: field scenario; 1,000: large sample size scenario) for five replicates per parameter set and fitted to eqn. (3) as described before (*n* = 60,000, of which 10,000 was burn-in).

## RESULTS

### Validating the inference methodology

We first validated the inference framework against data simulated with the system of ordinary differential equations described in eqn. (3), with an observation process that mimicked the field experiment and with parameters reflecting the range of values in the prior distributions.

*Large sample size scenario.*– In the large sample size scenario (five replicates with 1,000 released mosquitoes each), we accurately estimated the values of all parameters used in the simulations. All true parameter values fell within the 95% highest density interval (HDI) of the estimated posterior distributions (Figure 3). Most posterior medians approximated the true parameter well (Pearson *r* > 0.98), but somewhat less so for knockdown in the treated hut (Pearson *r* = 0.74). Posterior distributions were relatively wider for rate parameters associated with the treated hut (*x*_T_ and *k*_T_). Standard deviations of these parameters were a fraction (i.e., 11% and 12%) of their respective medians, whereas the s.d.:median ratio was below 3.5% for all other parameters. This reduced precision may be a consequence of the fact that rate parameters associated with huts other than the treated hut were informed by twice as much data as were the rate parameters associated with the treated hut, which derives from our assumption of shared parameters for huts a given distance from the treated hut (Figure 2). Gelman-Rubin statistics were below 1.1 for most simulation sets (average 1.04). When simulation sets resulted in parameters with GR statistics above 1.1, these were related to mosquito movement (*q*_i_, *r*_i_, and *p*_1_) and were most commonly associated with the untreated huts (Table S1). This indicates that those parameters may be among the most difficult to estimate.

**Figure 3:**
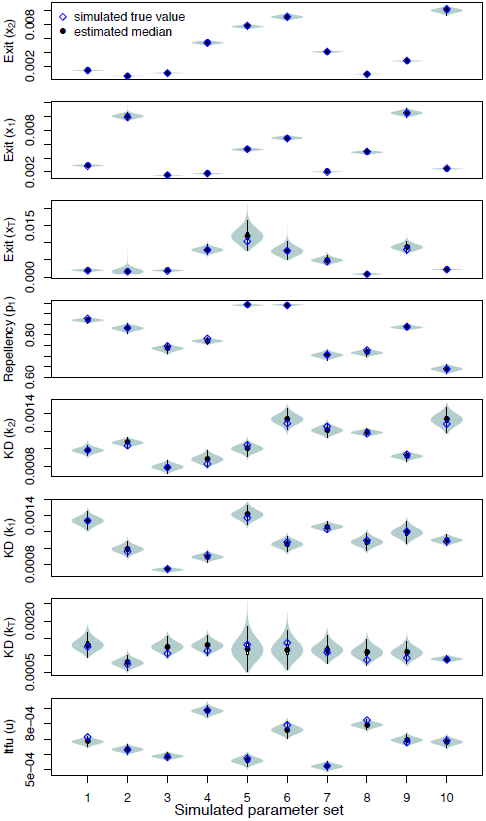
Estimated parameters from simulation experiments for five replicates of 1,000 released mosquitoes (large sample size scenarios) with the true value used in the simulation (blue diamonds) and the estimated median (black circles). The dashed gray line depicts *p*_1_ = 0.5, i.e., no repellency effect. Each estimate was based on five chains with distinct starting conditions. 60,000 MCMC iterations were performed inclusive of a burn-in period of 10,000.

***Field scenario***.–We also tested the performance of the inference framework on data simulated under conditions that closely resembled the conditions under which the experimental data were collected (Figure 4). All true parameter values fell within the 95% HDI, but the posterior medians were less consistent with the simulated values (*r* >0.8 for all but *x*_T_: 0.68; *k*_2_:0.03; *k*_T_: -0.29) than under the large sample size scenario (Figure 4). No systematic underestimation or overestimation was observed based on these simulations, suggesting that the additional discrepancy between simulated and inferred parameter values in the field scenario relative to the large sample size scenario was due to stochasticity associated with the smaller sample size in the field scenario (i.e., n = 25 vs n = 1,000). Gelman-Rubin statistics were, across all parameters and simulation sets, close to 1 (average GR 1.01) (Table S1).

**Figure 4:**
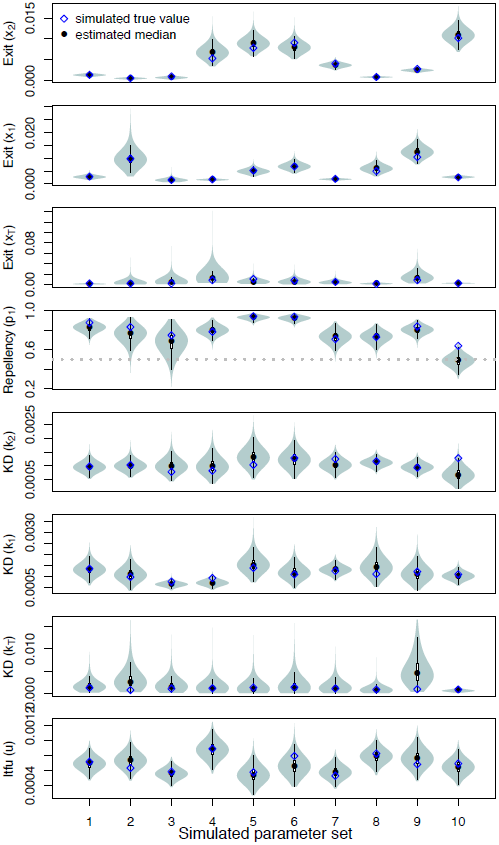
Estimated parameters from simulation experiments for five replicates of 25 released mosquitoes (field scenarios) with the true value (blue diamonds) and the estimated median (black circles). The dashed gray line depicts *p*_1_ = 0.5, i.e., no repellency effect. Each estimate is based on five chains with distinct starting conditions. 60,000 MCMC iterations were performed inclusive of a burn-in period of 10,000.

### Product effects on mosquito behavior

We first fitted the Markov chain model to the experimental hut data with all parameters allowed to vary. Strong correlations between *r*_*i*_*, q*_i_, and *p*_i_ indicated that these parameters were not identifiable given that a wide range of combinations of values of these parameters explained the data equally well (Figure S1). To resolve this identifiability issue, we fitted the exit rate *x*_*i*_ as a single composite parameter (*q*_*i*_*r*_*i*_). The rate of movement between huts is directly related to the exit rate; namely, it is a proportion (1-*r*_i_) of the overall movement rate out of a specific hut (*q*_*i*_). In doing so, we fixed the values of *r*_*i*_ at the medians of the posterior marginal density of the *r*_*i*_ corresponding to each hut that was obtained from the full parameter fit on the baseline data set (Figure S1). This reduced the amount of cross-correlation from Pearson’s *r* as high as 0.84 in the original to as low as 0.72 upon fixing *r*_*i*_. Most importantly, it markedly improved convergence from GR statistics as high as 1.38 (*q*_*2*_, low dosage) to as low as 1.00 for all parameters after fixing *r*_*i*_, indicating that other parameters became identifiable once this adjustment was made (Figure S7-Figure S9). Choosing either the 2.5^th^ or 97.5^th^ percentile of *r*_*i*_ instead did not affect this conclusion (Figure S5 and Figure S6). Acceptance rates for each chain tended to remain relatively constant following a burn-in period and varied across chains and parameters within the range of 21-54%.

*Exit and movement rates.*–Under baseline conditions (no chemical), exit rates (*x*_*i*_) from huts at different distances *i* from the treatment hut were relatively similar (medians for *x*_T_: 2.2×10^-3^, *x*_1_: 1.6×10^-3^, *x*_2_: 1.8×10^-3^) (Figure 5A-C). In subsequent treatment experiments, exit rates out of the treated hut were reduced relative to the baseline in response to both the low (RR = 0.70, HDI = 0.62-1.09) and the high transfluthrin dosage (RR = 0.70, HDI= 0.40-1.06), with no perceptible difference in the respective effects of the two dosages (Figure 5C). This effect carried over to the adjacent huts (*H*_1_) with exit rates lower than observed in the baseline experiment (low: RR = 0.79, HDI = 0.59-1.01; high: RR = 0.66, HDI = 0.50-0.87) (Figure 5B). In the huts furthest from the SR application (*H*_2_), the low dosage had no effect on exit rates relative to when no SR was applied (RR = 0.94, HDI = 0.72-1.18). To the contrary, the high dosage reduced exit rates (RR: 0.71, HDI:0.54-0.92) in all huts adjacent to the source of transfluthrin, including the furthest adjoining structures (Figure 5A). Given that the proportion of movement that was directed outdoors (*r*_*i*_) was held constant in this exercise, these results on exit rates (*q*_*i*_*r*_*i*_) are directly proportional to movement rates (*q*_i_).

**Figure 5:**
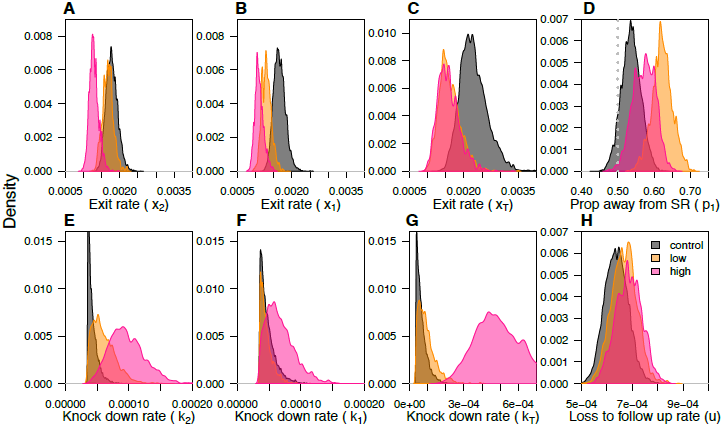
Posterior distributions of model parameters fitted to experimental data for the baseline (gray), low (orange) and high (pink) transfluthrin dosage for the treated hut (subscript T) and huts one or two removed from the treated hut (subscript 1 and 2, respectively). A-C) Rates at which mosquitoes exit the huts. D) Proportion of movement from *H*_1_ (hut directly adjacent to the treatment hut) away from the SR, where the dashed line indicates *p*_1_=0.5, i.e., no repellency effect. E-G) Knockdown rates. H) Loss to follow-up rates. The algorithm was run for 90,000 iterations inclusive of a burn-in period of 10,000.

**[fig5]**

*Repellency.*–In the baseline experiment, mosquitoes moved away from or towards the treated center hut (H_T_) with roughly equal probability (*p*_1_ = 0.54, HDI = 0.48-0.59), although with a possible slight preference for movement away from H_T_ (odds of moving away = 1.16, HDI = 0.92-1.41) (Figure 5D). In the experiment using low-dosage SR treatment, significant repellency from the treated center hut was observed (odds = 1.64, HDI = 1.30-2.09), with a median probability of moving away from this hut of 0.62 (HDI= 0.57-0.68) (Figure 5D). In the high-dosage treatment, repellency was still clear (odds = 1.35, HDI = 1.04-1.67), but the effect was somewhat smaller (*p*_1_ = 0.57, HDI = 0.52-0.63) (Figure 5D).

*Knockdown.*–Knockdown was a very rare event during baseline experiments (2/125 mosquitoes across all five replicates). As a consequence, estimates of knockdown rates in the baseline resembled the lower boundary of the prior distribution (medians for *H*_*T*_ = 5.8×10^-5^, *H*_1_ = 4.4×10^-5^, *H*_2_ = 4.0×10^-5^) (Figure 5E-5G). There was no effect of the low SR dosage on knockdown rates relative to the baseline, both in the treated hut *H*_*T*_ (RR = 1.39, HDI = 0.26-3.84) (Figure 5G) and in the *H*_1_ huts directly adjacent (RR = 1.00, HDI = 0.45-1.76) (Figure 5F). In the *H*_2_ huts furthest away from the treatment, a somewhat increased knockdown rate was observed in response to the low dosage relative to the baseline (RR = 1.37, HDI = 0.64-2.46) (Figure 5E). Knockdown rates in the high-dosage scenario were elevated in all huts, in particular in the *H*_*T*_ treatment huts (RR = 8.37, HDI = 2.11-17.35) (Figure 5G) but also in the *H*_1_ and *H*_2_ huts (H_1_: RR = 1.39, HDI = 0.52-2.69; H_2_: RR = 2.22, HDI = 0.96-3.86) (Figure 5E and 5F).

*Loss to follow-up.*–Rates of loss to follow-up were similar across the baseline and two SR treatment experiments, although there was a signal for a small increase in these rates with increasing dosage (low: 5%, high: 8%) (Figure 5H). In comparing posterior samples across dosages, a signal for a positive dose-response relationship (i.e., *u*(high) > *u*(low) > *u*(baseline)) was confirmed in 61% of samples from the posterior.

*Time spent in a hut*.–The total amount of time a mosquito spent in each hut results from the composite of treatment effects. By running simulations of the system of ordinary differential equations (eqn. (3)) with the estimated posterior parameter values, we derived a posterior estimate of the proportion of the time a mosquito spent in each hut relative to the total time a mosquito was in the hut system (i.e., before exit, kd, or ltfu). This proportion was found to be similar but slightly reduced for the treated hut *H*_T_ relative to the baseline scenario in either treatment scenario (Figure 6E) and without any effect in the downstream huts *H*_1_ and *H*_2_ (Figure 6A and 6C). However, when considering the total duration of the experimental day, the proportion of time spent in the adjacent, downstream huts *H*_1_ and *H*_2_ was higher during experiments using both low and high SR dosage than during baseline (Figure 6B and 6D). This was a result of reduced exit rates and thus an overall increase in time spent in the hut system as a whole (Figure 5A and 5B).

**Figure 6:**
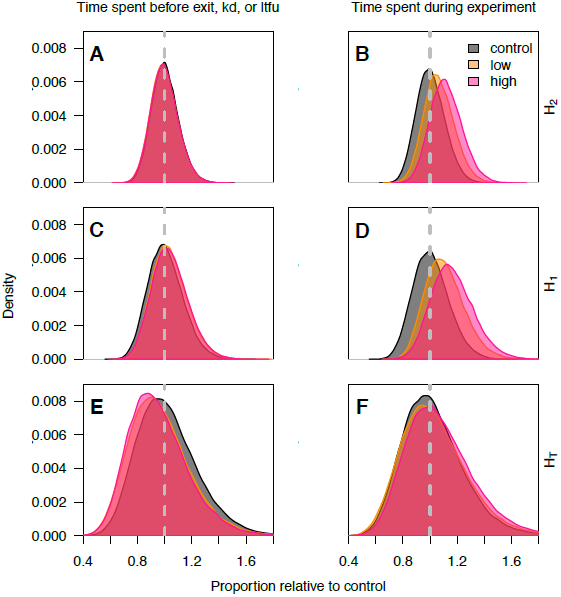
Distributions of time mosquitoes spent in each hut relative to baseline (gray), low (orange), and high (pink) transfluthrin dosage for huts two (A and B) or one (C and D)removed from the treated hut (E and F). The left column signifies the proportion of time spent in each hut before having experienced an event (A, C, and E), where kd is knockdown and ltfu is loss to follow-up. The right column signifies the proportion of the total experiment time spent in each hut relative to the baseline (B, D, F).

## DISCUSSION

Alternative *Ae. aegypti* vector control strategies are currently being evaluated to address challenges related to dengue transmission expansion [2]. Spatial repellent (SR) products, which release volatile chemicals into treated spaces to interrupt host-vector contact, are among these [14]. One challenge for evaluating the efficacy of SRs, and other products that may include non-lethal outcomes, is characterizing the multifaceted effects of a given product on mosquito behavior under field conditions, which is critical for guiding product formulation and implementation under operational conditions.

To quantify concurrent and downstream *Ae. aegypti* behavioral effects of a transfluthrin-based experimental spatial repellent (SR) product, we used a continuous-time Markov chain model informed by experimental data under a Bayesian inference framework. Examination of posterior estimates of model parameters showed that test mosquitoes were deterred from entering the experimental hut where the product was located and that this effect was stronger for the lower dosage SR. Posterior estimates of model parameters also indicated lower movement rates out of the treatment hut (either to a neighboring hut or out of the hut system) under both treatments, presumably due to confusion effects at both dosages. Under the higher SR dosage, the reduced movement effect was noticeable in adjacent untreated huts as far as two huts away from the SR application. Similarly strong downstream effects were observed on knockdown rates, which were markedly increased in all huts in the presence of the higher SR dosage, yet little effect on knockdown was observed at the lower dosage.

We validated our inference method by demonstrating its ability to accurately estimate the model’s parameters given simulated data. This assessment was conditional, however, on the assumption that the model is a realistic representation of reality. Some of the known limiting assumptions of our analysis include (1) effects that depend on distance from the treated hut rather than on each hut individually, (2) equal loss to follow-up across huts, and (3) time-invariant parameters. Of these, the first may be most problematic when considering that air flow within the hut system could result in asymmetric effects of transfluthrin dispersion to huts of the same distance from the treated hut but on different sides of it [44]. In principle, it would be possible to account for such factors in future studies by measuring air flow and incorporating its effect on the data through an appropriate modification of the model. For example, repellency (*p*_*1*_) could be allowed to vary across huts and treated as a function of readings from a wind gauge. Posterior estimates of the parameters governing the relationship between wind and *p*_*1*_ would then allow for inferences about the repellency of the product under varying airflow conditions and beyond those observed in the current experiment. Planning for required sample sizes and sampling schemes for such experiments would benefit from our model and results by using our posterior distributions to inform prior distributions in those future studies [45].

Repellency and increased knockdown reduced the overall time that mosquitoes spent in a transient, non-terminal state in the experimental hut with the SR application, whereas decreased movement rates have the potential to offset this effect. The result of the product’s impact on the time mosquitoes spent in the treated hut indicates potential for such a product to limit human-vector contact (and thereby reduce the probability of pathogen transmission) in the treated hut. However, it is uncertain to what extent host-seeking and blood-feeding behaviors of these mosquitoes exposed to the SR may have been affected in the current study. Other studies using similar volatile products have shown these effects to also be associated with reduced rates of human landing [13]. The inclusion of blood-feeding metrics in experiments with volatile pyrethroid products using anophelines under field conditions [46] and against the topical repellent DEET using *Ae. aegypti* in the laboratory [47] have been valuable in establishing expectations of such synergistic chemical effects.

The effect of SR products on untreated neighboring premises has been a consistent and critical question to the public health value of these products [14, 48]. Three aspects of our results suggest that the risk of diversion of mosquitoes from a treated area to adjacent untreated areas may be limited for the formulation used in our experiments. First, SR exposure reduced movement rates between huts. Second, the time spent in untreated huts relative to the treated hut was unaffected by the presence of the SR relative to the baseline, once reduced exit rates were accounted for. Third, there was a marked increase in knockdown in untreated huts at the high SR dosage. At the same time, there was also a marked reduction in exit rates out of untreated huts, which resulted in prolonged time spent in adjacent huts. Evaluating the overall potential for diversion based on these effects will require pairing experimental results such as ours with new theory that is capable of translating this range of behavioral effects into estimates of their epidemiological consequences [15, 49, 50].

Under our experimental design, we cannot distinguish between downstream effects caused by volatile particles dispersed into untreated huts or by a residual, post-exposure effect of transfluthrin on mosquitoes that are exposed in the treated hut but exit or are knocked down elsewhere. The reduced repellency effect observed at high relative to low dosage may not result from reduced sensitivity of mosquitoes to this effect per se, but may instead be a result of saturation of all experimental huts with the volatile chemical. Indeed, correlations between air sampling measurements and mosquito behavior responses have been explored in previous studies using spatial repellents but with limited success due to limits of chemical detection and quantification [51-53]. Combining air chemistry inferences of specific active ingredients (i.e., vapor pressure, particle weight) with environmental data (i.e., air current, flow rate) into our new inference framework is therefore warranted and could enable quantification of the extent to which downstream effects result from movement of the volatile chemical or movement of exposed mosquitoes with lingering post-exposure effects. The latter possibility has been indicated in other studies to have potential for innovative applications of SRs [54].

## CONCLUSIONS

The need for development and efficient testing of new vector control products and innovative formulations of existing tools is evident [2, 55]. The framework we introduce here provides a flexible tool to estimate a product’s effects on mosquito vector movement in a quantitative and probabilistic fashion, without the need for expensive, technologically advanced mosquito tracking devices. In addition, context-specific diversion (movement to an untreated space) and downstream (adjacent spaces) effects can be estimated at early stages of product development under scenarios similar to operational settings for which the product is intended to be used. The value of our inferential framework is best highlighted when considering the need for a cheap, cost-effective, efficient, and accurate methodology for the characterization and down-selection of novel vector control products (or combinations thereof). This not only supports early optimization but also the establishment of target product profiles to identify those products under development that are worth advancing to costlier epidemiological trials. More specifically to SRs, our results strengthen expectations of proof of concept of spatial repellents regarding public health value required for WHO recommendations [56].

## LIST OF ABBREVIATIONS

FAR: field application rate
GR: Gelman-Rubin
HDI: highest density interval
IRS: indoor residual spraying
ITN: insecticide treated net
ltfu: loss to follow-up
MCMC: Markov chain Monte Carlo
MRR: mark-release-recapture
RR: relative risk
SR: spatial repellent

## DECLARATIONS

### Ethics approval and consent to participate

Not applicable

### Consent for publication

Not applicable

### Availability of data and material

Code used to perform the analyses are available on Github.com (https://github.com/quirine/ExperimentalHuts) with output data deposited on OSF.io (https://osf.io/xtmy7/) as part of project (https://osf.io/5hcpf/).

### Authors’ contributions

FCL, HM, ACM, JPG, and NLA designed the hut experiments. FCL and HM performed the experiments. QAtB and TAP developed the modeling and simulation framework. QAtB performed the analyses. QAtB, JPG, NLA, and TAP interpreted the results. QAtB, NLA, and TAP wrote the manuscript. All authors read, provided feedback, and approved the final manuscript.

### Acknowledgements

We express our gratitude to Carlos Ique, Director of the Institute Veterinario de Investigaciones (IVITA), Iquitos, Peru for the use of land for experimental hut studies. Thanks to Roxanne Burrus, Kirk Mundal, and Victor Lopez (NAMRU-6) for their logistic support during the experiments and to Neil Lobo for valuable insights. Special thanks to Edwin Requena, Hugo Jaba, and to the team of mosquito collectors for their dedicated effort and to reviewers for valuable comments.

### Disclaimer

The views expressed in this work are those of the authors and do not reflect the official policy or position of the Department of the Navy, Department of Defense, or U.S. Government.

### Copyright statement

FCL and ACM are employees of the U.S. Government. This work was prepared as part of her official duties (PJTNMRCD 0.24). Title 17 U.S.C. § 105 provides that ‘Copyright protection under this title is not available for any work of the United States Government’. Title 17 U.S.C. § 101 defines a U.S. Government work as a work prepared by a military service member or employee of the U.S. Government as part of that person’s official duties.

## SUPPORTING TABLE AND FIGURE LEGENDS

Table S1: Average Gelman-Rubin statistics across simulated data sets (median and the upper bound of the 95% confidence interval).

Figure S1: Correlations between parameter posteriors of model fit on baseline scenario with all parameters estimated. Marginal posteriors are depicted on the diagonals. The numbers on the right of the diagonal depict the correlation coefficients for each side by side comparison.

Figure S2: Correlations between parameter posteriors of model fit on baseline scenario with ri fixed. Marginal posteriors are depicted on the diagonals. The numbers on the right of the diagonal depict the correlation coefficients for each side by side comparison.

Figure S3: Correlations between parameter posteriors of model fit on low dosage scenario with ri fixed. Marginal posteriors are depicted on the diagonals. The numbers on the right of the diagonal depict the correlation coefficients for each side by side comparison.

Figure S4: Correlations between parameter posteriors of model fit on high dosage scenario with ri fixed. Marginal posteriors are depicted on the diagonals. The numbers on the right of the diagonal depict the correlation coefficients for each side by side comparison.

Figure S5: Posterior distributions of model parameters fitted to experimental data while fixing the values of ri at the 2.5th percentile of the posterior from the full parameter fit to the baseline data. Posteriors are shown for the baseline (gray), low dosage (orange) and high dosage (pink) for the SR-hut (subscript 0) and huts 2 or 1 removed (subscript 2 and 1 respectively). A-C) rates at which mosquitoes exit the huts D) proportion of movement from H1 (hut directly adjacent to the treatment hut) away from the SR-product. E-G) knockdown rates, and H) loss to follow-up rates. The algorithm was run for 25,000 iterations with a burn-in period of 10,000.

Figure S6: Posterior distributions of model parameters fitted to experimental data while fixing the values of ri at the 97.5th percentile of the posterior from the full parameter fit to the baseline data. Posteriors are shown for the baseline (gray), low dosage (orange) and high dosage (pink) for the SR-hut (subscript 0) and huts 2 or 1 removed (subscript 2 and 1 respectively). A-C) rates at which mosquitoes exit the huts D) proportion of movement from H1 (hut directly adjacent to the treatment hut) away from the SR-product. E-G) knockdown rates, and H) loss to follow-up rates. The algorithm was run for 25,000 iterations with a burn-in period of 10,000.

Figure S7: Gelman-Rubin convergence diagnostics by iteration for the baseline scenario. Figure S8: Gelman-Rubin convergence diagnostics by iteration for the low dosage scenario. Figure S9: Gelman-Rubin convergence diagnostics by iteration for the high dosage scenario.

Figure S10: Trace plots for the baseline scenario. Figure S11: Traceplots for the low dosage scenario. Figure S12: Traceplots for the high dosage scenario.

